# Modeling bee movement shows how a perceptual masking effect can influence flower discovery, foraging efficiency and pollination

**DOI:** 10.1101/2022.09.12.507525

**Authors:** Ana Morán, Mathieu Lihoreau, Alfonso Pérez Escudero, Jacques Gautrais

**Affiliations:** Centre de Recherches sur la Cognition Animale (CRCA), Centre de Biologie Intégrative (CBI), Université de Toulouse, CNRS, UPS, 118 route de Narbonne, Bat 4R4, 31062 Toulouse Cedex 9, France

**Keywords:** central place foraging, bee, model simulations, persistent turning walker

## Abstract

Understanding how pollinators move across space is key to understanding plant mating patterns. Bees are typically assumed to search for flowers randomly or using simple movement rules, so that the probability of discovering a flower should primarily depend on its distance to the nest. However, experimental work shows this is not always the case. Here, we explored the influence of flower size and density on their probability of being discovered by bees by developing a movement model of central place foraging bees, based on experimental data collected on bumblebees. Our model produces realistic bee trajectories by taking into account the autocorrelation of the bee’s angular speed, the attraction to the nest, and a gaussian noise. Simulations revealed a « masking effect » that reduces the detection of flowers close to another, which may have critical consequences for pollination and foraging success. At the plant level, flowers distant to the nest were more often visited in low density environments, suggesting lower probabilities of pollination at high densities. At the bee colony level, foragers found more flowers when they were small and at medium densities, suggesting that there is an optimal flower size and density at which collective foraging efficiency is optimized. Our results indicate that the processes of search and discovery of resources are potentially more complex than usually assumed, and question the importance of resource distribution and abundance on plant-pollinator interactions.

**Author’s summary:** Understanding how pollinators move in space is key to understanding plant reproduction, which in turn shapes entire ecosystems. Most current models assume simple movement rules that predict that flowers are more likely to be visited—and hence pollinated—the closer they are to the pollinators’ nest. Here we developed an explicit movement model that incorporates realistic features of bumblebees, including their flight characteristics and their tendency to return regularly to the nest, and calibrated it with experimental data collected in naturalistic conditions. This model revealed that the probability to visit a flower does not only depend on its position, but also on the position of other flowers that may mask it from the forager. This masking effect means that pollination efficiency depends on the density and spatial arrangement of flowers around the pollinator’s nest, often in counter-intuitive ways. Taking these effects into account will be key for improving precision pollination and pollinator conservation.

## Introduction

Pollinators, such as bees, flies, butterflies, but also bats and birds, mediate a key ecosystemic service on which most terrestrial plants and animals, including us humans, rely on. When foraging for nectar, animals transfer pollen between flowers, which facilitates plant reproduction. Understanding how pollinators move, find and choose flowers is thus a key challenge of pollination ecology (1). In particular, this may help predict and act on complex pollination processes in a context of a looming global pollination crisis, when food demand increases and pollinators decline (2,3).

Pollinators have long been assumed to move randomly (4–7) or use hard wired movement rules such as visiting the nearest unvisited flower (8), exploiting flower patches in straight line movements (9), navigating inflorescences from bottom to top flowers (10), or using win-stay lose-leave strategies (11). Therefore, pollination models relying on these observations typically predict diffusive movements in every direction (12). However, recent behavioral research shows this is not true when animals forage across large spatial scales (13). In particular, studies using radars to monitor the long distance flight paths of bees foraging in the field demonstrate foragers learn features of their environment to navigate across landscapes and to return to known feeding locations (14,15). This enables them to develop shortcuts between food sources (16) and build efficient multi-location routes (also called “traplines”) minimizing overall travel distances (17,18). These routes are re-adjusted each time a food source is depleted and new ones are discovered (19).

How bees learn such foraging routes has been modelled using algorithms implementing spatial learning and memory (20–22). While this has greatly advanced our understanding of bee exploitative movements patterns, none of these models have looked at search behaviors, either assuming insects already know the locations of all available feeding sources in their environment or discover them according to fixed probabilistic laws (i.e. the probability to discover a flower at a given location is proportional to 1/*L*^2^ where *L* represents the distance to that flower (20–22)). However, experimental data indicate that this is not the case. Firstly, bees, like many pollinators, are central place foragers (i.e., every foraging trip starts and ends at the nest site (23)). This means that their range of action is limited. Recordings of bee search flights, although scarce, show how individuals tend to make loops centered at the nest when exploring a new environment and look for flowers (14,24). These looping movements are not compatible with the assumption that bees make diffusive random walks or Lévy flights (25,26). Secondly, the spatial structure of the foraging environment itself may also greatly influence flower discovery by bees. In particular, the probability of finding a flower heavily depends on the location of the flower visited just before, ultimately influencing the direction and geometry of the routes developed by individuals (17–20,27,28). Since bees are more attracted by larger flowers than by smaller ones (29), this suggests that small isolated flowers could be missed if they are located next to a larger patch. Ultimately, this potential « masking effect » on the probability to visit specific flowers depending on the presence of other flowers around could have important consequences for bee foraging success, for instance by precluding the discovery of some highly rewarding flowers that are isolated or further away from the nest. This could also dramatically influence plant pollination, if bees are spatially constrained to single flower patches and plant outcrossing is limited.

Here we developed a model of bee search movement simulating the tendency of bees to make loops around their nest and examined the influence of these looping flights on the probability for bees to discover flowers in environments defined by resources of various sizes and abundances. We hypothesized that considering looping movements characteristic of bee exploratory flights would result in strikingly different predictions than the typical diffusive random walk movements. We also described perceptual masking effects by which the probability of finding given flowers is affected by the presence of others, and analyzed their consequences on discovery rates, bee foraging efficiency and plant pollination success.

## The model

### Description

Our model is an extension of the Persistent Turning Walker (PTW) developed by Gautrais et al. (30) to model fish movements in a tank. For the sake of simplicity, here we modelled bee movements in 2D, neglecting altitude. We assumed that bees fly at constant speed and with varying angular speed, *ω*(*t*), which is governed by

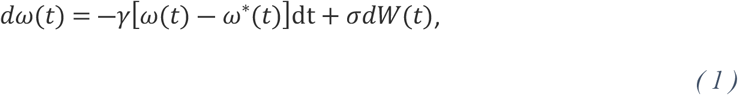

where *γ* is an auto-correlation coefficient and *σ*d*W*(*t*) introduces a gaussian noise, governed by a Wiener process (31). The two terms of Equation 1 have opposing effects: The first term pushes the angular speed towards a target angular speed, *ω*(*t*), with a strength controlled by the auto-correlation coefficient *γ.* The second term introduces noise in the angular speed making bees change direction. Therefore, high values of *γ* lead to smoother and more predictable trajectories. Setting *ω*\*(*t*) = 0 leads to a trajectory with no preferred direction, whose angular speed changes smoothly around zero. This is the simplest condition, resembling a diffusive process in which the animal moves aimlessly and gets further and further from its initial position as time goes by (32–37).

We modelled central place foraging by adding an attraction component to the model in order to make bees return to the nest after a certain amount of time. To implement the return to the nest we assumed that bees can locate the direction of their nest at any time using path integration (i.e. navigational mechanism by which insects continuously keep track of their current position relative to their nest position (38)), and define a homing vector, 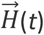 that points towards the nest (13). Then, we assumed that the bee tries to target the angular speed that will align its trajectory with the homing vector, so we modeled the target angular speed as

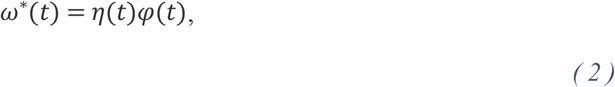

where *φ*(*t*) is the angle between the bee’s heading 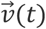 and the homing vector 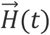 (Fig. 1A), and *η*(*t*) is the attraction strength that controls a switch between the exploration and return phases: During the initial exploration phase we make *η*(*t*) = 0, so that bees explore randomly and distance themselves from the nest, while during the return phase we make *η*(*t*) = *η\** > 0, so that the bee has a tendency to turn towards the nest. We assumed that bees switch instantaneously between the exploration and return phases, so

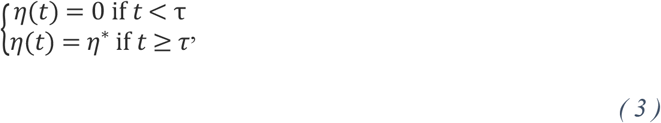

where *τ* is the time at which the switch happens (Fig. 1B). This switch may happen at any time, with a constant probability per unit time, *p*_return_. This means that the switching times are exponentially distributed, with an average time of approximately 1/*p*_return_.

**Fig 1.**
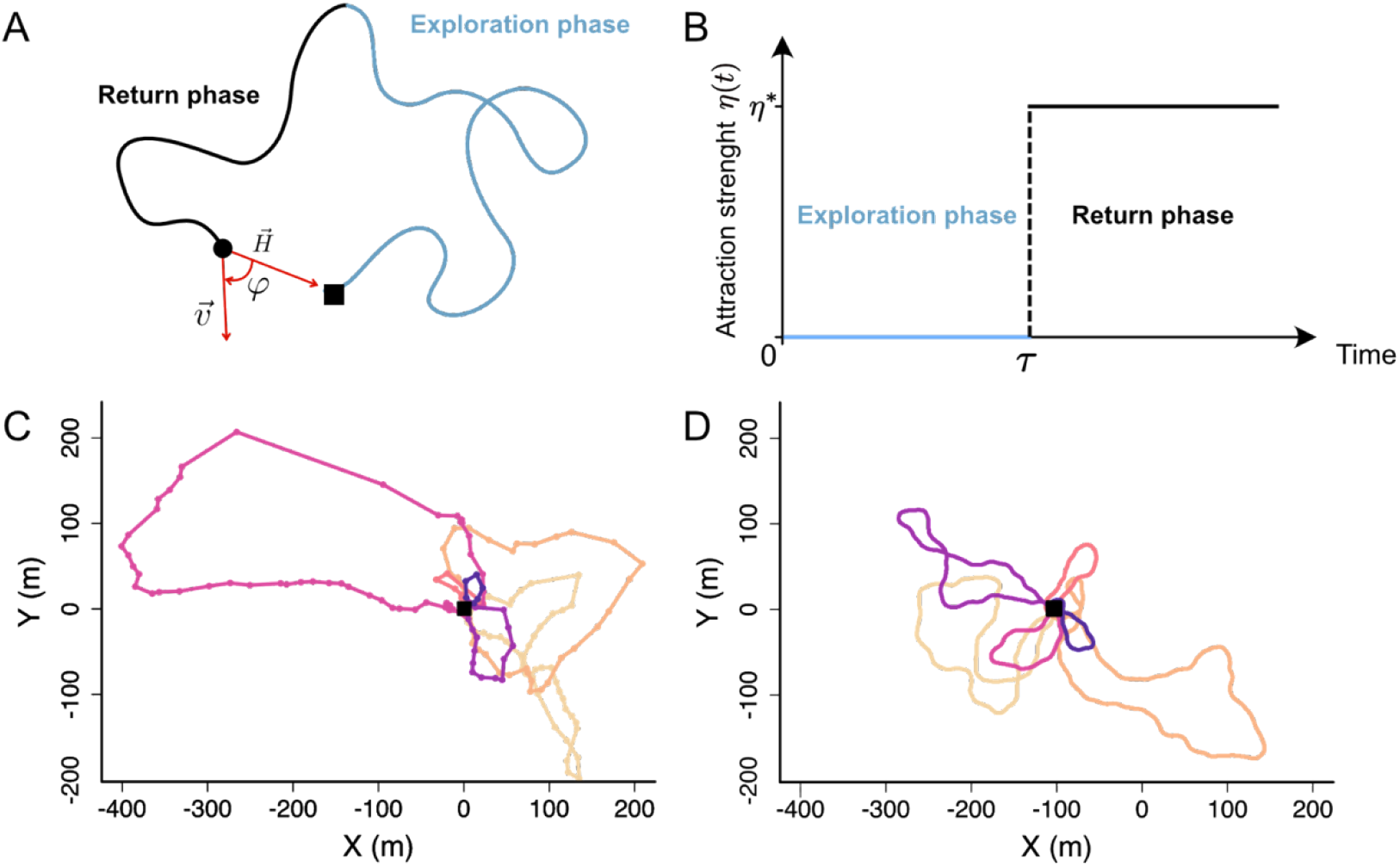
Illustration of the model. **(A)** Example of theoretical trajectory. Blue line: Trajectory during the exploration phase. Black line: Trajectory during the return phase. Black circle: bee. Black square: nest. H is the homing vector pointing towards the nest. 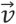 is the heading of the bee. *φ* is the angle between 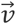 and 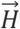. **(B)** Evolution of the return strength (*η*) over time. At time = τ, η switches from 0 (no attraction) to *η*\*. **(C)** Example of an experimental trajectory (39). Each dot represents the position of a bee recorded by a harmonic radar approximately every 3s. Different colors represent different loops around the nest. The sequential order of the loops is represented by the color gradient where the first loops have lightest colors (yellow to purple). **(D)** Same as C, but for a simulated trajectory with parameters *γ* = 1.0 *s*^-1^, *σ* = 0.37 rad/s^1/2^, *p*_return_ = 1/30 *s*^-1^ and *η*\* = 0.2 *s*^-1^.

The model therefore has four main parameters: The auto-correlation (γ) and the randomness (σ) control the characteristics of the flight, while the probability to return (*p*_return_) and the strength of the attraction component (*η*\*) control the duration of each exploration trip. Here we have described the continuous version of the model, but to implement it numerically we discretized it in finite time steps (see Methods).

### Calibration with experimental data

In principle, our model can describe search movements of any central place forager. Here we explored its properties focusing on a model species for which we had access to high-quality experimental data: the buff-tailed bumblebee *Bombus terrestris.* We used the dataset of Pasquaretta et al. (39) in which the authors used a harmonic radar to track 2D exploratory flight trajectories of bees in the field. Bees carrying a transponder were released from a colony nest box located in the middle of a large and flat open field, and performed exploration flights without any spatial limitation. The radar recorded the location of the bees every 3.3s over a distance of ca. 800 m (Fig 1C). In these experiments the bees were tested until they found artificial flowers randomly scattered in the field. We used 32 radar tracks for 18 bees.

To quantify the experimental trajectories, we first divided tracks into “loops”, each loop being a segment of trajectory that starts and ends in the nest (Fig 1C). This extraction yielded 207 loops. We then computed four observables for each loop (Fig 2):

- Loop length: Total length of the trajectory for a given loop (Fig. 2A).
- Loop extension: Maximum distance between the bee and the nest for a given loop (Fig. 2B).
- Number of intersections: Number of times the loop intersects with itself (Fig. 2C).
- Number of re-departures, where a re-departure is defined as three consecutive positions such that the second position is closer to the nest than the first one, but the third is again further away than the second. These events indicate instances in which the bee seemed to be returning towards the nest and turned back (Fig. 4D).

**Fig 3.**
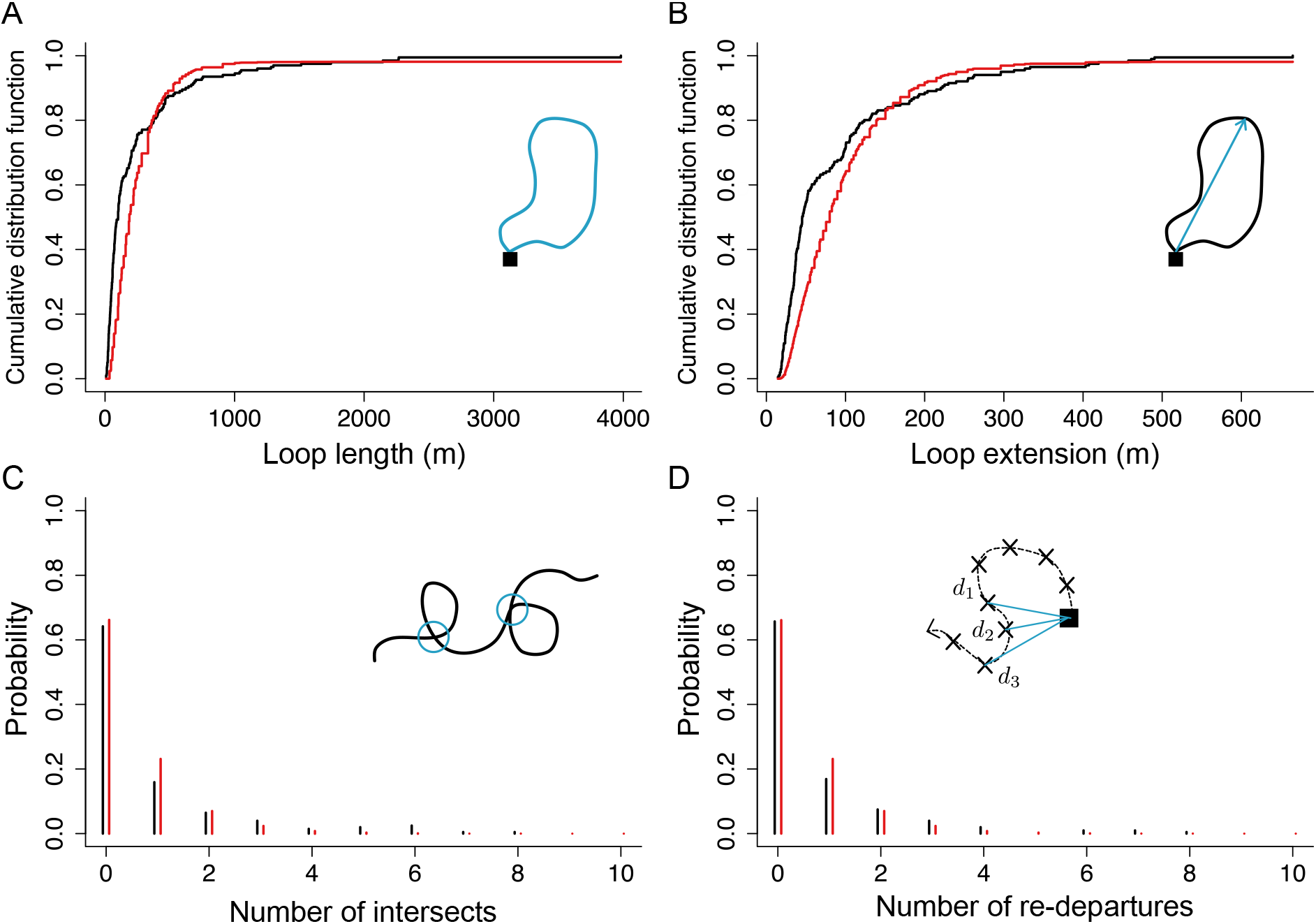
Distributions of the four observables, for experimental and simulated data. Black lines: experimental data. Red lines: model predictions using the optimal *γ* = 1.0 *s*^-1^, *σ* = 0.37 rad/s * s^1/2^, *p*_return_ = 1/30 *s*^-1^ and *η*\* = 0.2 *s*^-1^. Insets: Schematic of each observable. **(A)** Cumulative distribution function of loop lengths for our full dataset. **(B)** Same as A, but for the loops extension. **(C)** Probability distribution of the number of trajectories intersects per 100m traveled. **(D)** Same as C, but for the number of re-departures per 100 m traveled.

We extracted these four parameters from each loop and found substantial variability in all of them (Fig. 3, black lines). We then used this information to find the optimal model parameters, aiming to describe not only the average value of each observable, but also their variability. To do so, we performed simulations covering exhaustively all relevant combinations of our four parameters. For each combination of parameters, we simulated 1000 loops, extracted the distributions for the four observables, and chose the parameter combination that best approximated the experimental distributions for the four observables (see Methods). This procedure resulted in the optimal parameters *γ* = 1.0 *s*^-1^, *σ* = 0.37 rad/s * s^1/2^, *p*_return_ = 1/30 *s*^-1^ and *η\** = 0.2 *s*^-1^, which give a good approximation to the experimental distributions of observables (Fig. 2, red lines), and trajectories that qualitatively resemble the experimental ones (Fig. 1D)

**Fig 3.**
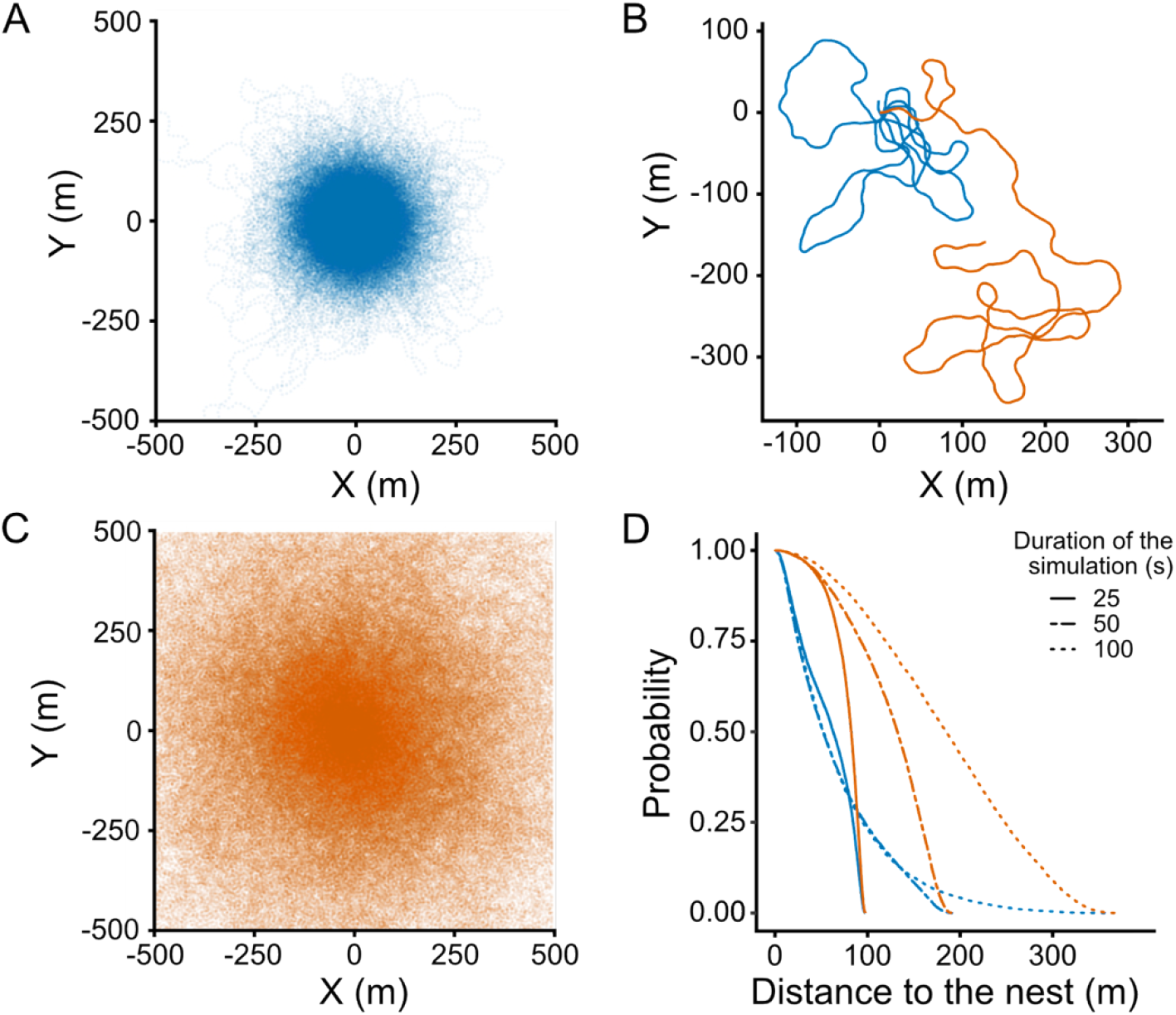
Probability of presence of a bee around the nest. **(A)** Overlay of 1000 trajectories with attraction to the nest (*η*\*= 0.2 *s*^-1^) simulated during 900 s. **(B)** Same as A but without attraction to the nest (*η\** = 0) (**C)** Probability to find a bee below a given distance to the nest (i.e., inverse cumulative probability distribution for the distance to the nest) after different amounts of time. **(D)** Example trajectories with and without attraction, simulated during 500 s. The nest is located at (0,0). Blue: model with attraction. Orange: model without attraction.

## Model predictions

### Attraction to the nest limits the exploration range of bees

An unrealistic feature of existing diffusive models is their long-term behavior: If given enough time, the forager reaches extremely far distances with respect to the nest, never returning to it. To measure the impact of central place foraging on bee exploration range, we compared our model with attraction to the nest to an alternative one in which the attraction is absent (i.e., making *η*\* = 0 in Equation 3). We then simulated 1000 trajectories with each model for different amounts of time, and studied how the distribution of bees around the nest changes over time.

This analysis revealed two main effects of the attraction component. First, it retains bees tightly localized around the nest (~250m) (Fig. 3A-B, blue). Second, it makes the distribution of bees stationary: In a model without attraction, bees constantly wander away from the nest, and their distribution depends on how much time we allow for the bee to explore, becoming wider as time goes by (Fig. 3B-C, orange). In contrast, the attraction component makes the forager return to the nest periodically, so the distribution remains constant once the forager has had enough time to perform more than one loop on average (Fig. 3D, blue)

### Distant flowers are more often visited in low-density environments

A key consequence of bee movement is its influence on plant reproduction through pollen dissemination. We estimated the probabilities of flowers to be discovered (and thus pollinated) by bees in a field characterized by a random and uniform distribution of flowers, an average density of 1.3 10^-4^ flowers/m2 and a diameter of 70cm (for the sake of simplicity here a “flower” is equivalent to a feeding location, which may be a single flower or a plant containing several ones). We assumed that a flower was visited whenever its distance to the bee’s trajectory was below a threshold, given by the bee’s visual perception range (see Methods). We focused on vision rather than olfaction because vision is the main sense that bees use to accurately navigate the last meters towards a particular flower, while olfaction is used at a broader spatial scale (23). Using these conditions, we simulated 1000 foraging trips, each of them lasting 900 seconds, and for each flower we computed the probability to be found in a given trip (i.e., the proportion of simulations in which the trajectory overlaps with the flower’s area of attraction). This probability falls exponentially with the distance between the flower and the nest (Fig. 4A, red line).

**Fig 4.**
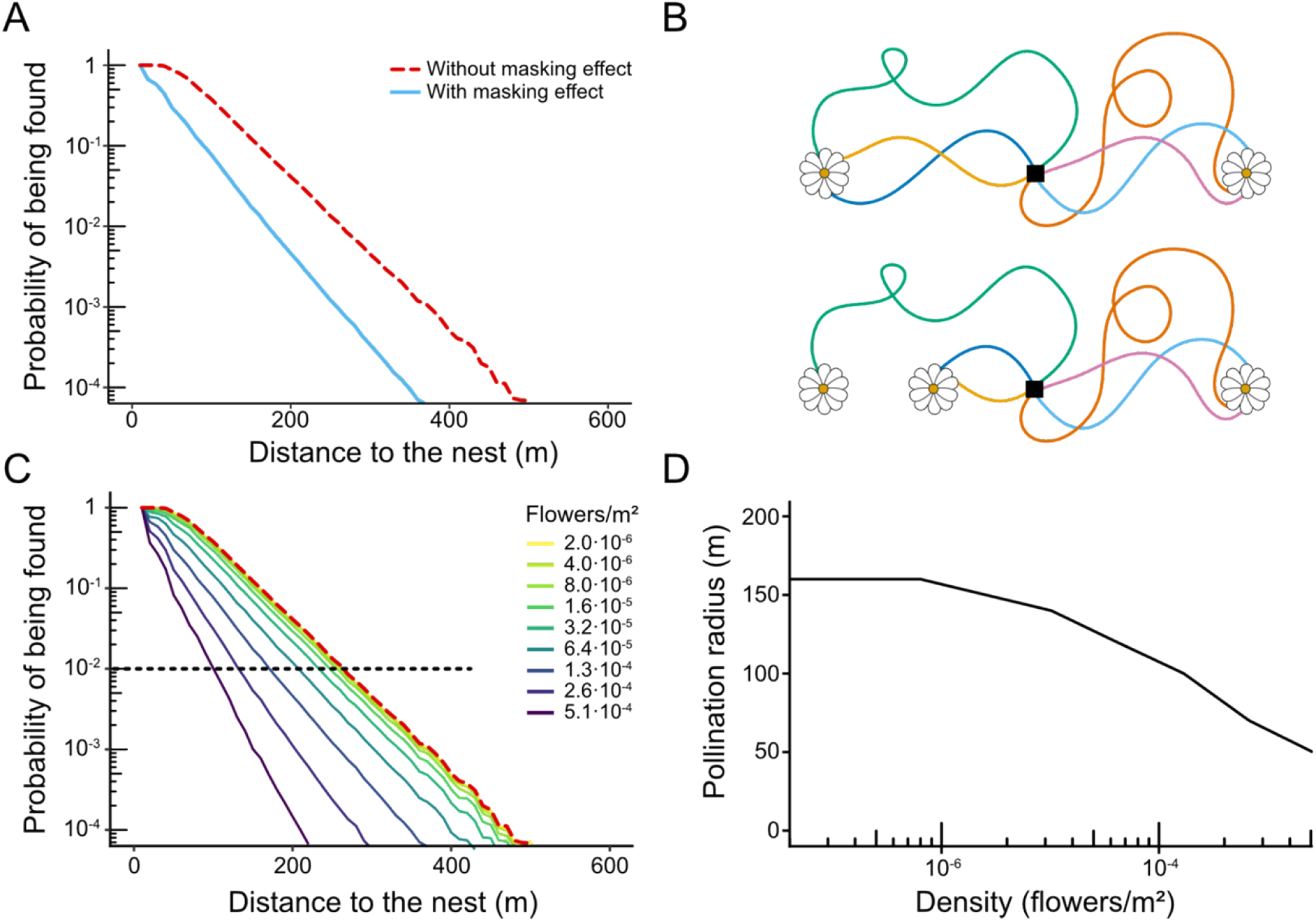
Predicted pollination efficiency. **(A) Probability that a flower is found as a function of its distance to the nest.** We simulated exploration trips in a field of uniformly distributed flowers with density 1.3 10^-4^ flowers/m^2^ and flower size 70 cm. For each flower, we computed the probability that it was found in each exploration trip, and we show this probability as a function of the distance between the flower and the nest. Results computed over 6000 simulated trips of 900s in 10000 environments for each density. Red line: Probability calculated without taking into account the masking effect. Blue line: Probability calculated taking into account the masking effect (i.e., only counting the first flower that was discovered in each trip). **(B) Illustration of the masking effect.** The probability of finding a flower depends on the presence of other flowers. In a scenario where there are just 2 flowers equidistant to the nest, both flowers should be visited equally (top). However, if another flower is added, it can capture visits that would otherwise visit one of the original flowers (bottom). Black square: nest. **(C)** Same as (A), but for different flower densities. Red dotted line: Probability calculated without taking into account the masking effect. This probability is independent of the density of flowers. Solid lines: Probability calculated taking the masking effect into account. Black dotted line: threshold probability at which we consider an area that has a high probability of being pollinated. **(D)** Radius of the area around the nest that has a high probability of being pollinated (i.e., where the probability that flowers are discovered is above 10^-2^) as a function of flower density.

However, this simulation does not take into account the fact that the probability of visiting a flower does not only depend on its distance to the nest, but can also be influenced by the presence of other flowers around. This dependence exists because a bee that finds a flower does not continue its trajectory, but will rather stop to collect nectar. Once nectar collection is over, the bee may continue exploring, but after visiting a few flowers the bee returns to the nest to unload its crop. For example, in a scenario where there are just 2 flowers equidistant to the nest, both flowers should be visited equally. However, if another flower is added, it can capture visits that would otherwise visit one of the original flowers, reducing the probability that it’s discovered (Fig. 4B). We call this “the masking effect” (Figure 4B).

In order to model this masking effect in a simple way, we assumed that each bee returns to the nest after discovering a single flower. The first qualitative consequence of the masking effect is to reduce the probability that flowers distant to the nest are discovered (Fig. 4A, blue). The second consequence is that it introduces a dependence of flower density on pollination efficiency. In the absence of masking, only two factors determine the probability that a flower is discovered: its size (which determines the distance from which it can be perceived) and its distance to the nest. In contrast, when masking is taken into account, the number of visits also depends on the overall density of flowers in the environment, falling more sharply with distance when this density is higher (Fig. 4C).

This dependence with flower density means that the area around the nest where flowers have a high probability of being pollinated depends on flower density. To estimate the size of this area, we set a threshold at a probability of 10^-2^ per trip (black dotted line in Fig. 4C), and computed the “pollination radius” as the distance at which flowers’ probability of being discovered remains above this threshold, assuming than one visit is enough for pollinating a plant. At low flower densities the pollination radius reaches 160 meters, and is limited by the bees’ exploration range (i.e., their tendency to return to the nest after a certain time, even if no flowers have been found; compare this radius with the distribution in Fig. 3C). Due to the masking effect, the pollination radius decreases as flower density increases (Fig. 4D).

### Bees find more flowers at intermediate densities

We then explored potential influences of the masking effect on the foraging success of the bees. This success depends on the number of flowers discovered collectively by all the bees of a colony, because a flower discovered and exploited by a bee will be at least partially depleted, giving little additional benefit to later visitors. For this reason, what counts is not the total number of visits that bees perform, but rather the total number of different flowers discovered by the colony. One would expect that higher flower density would make it easier for the colony to discover a higher number of flowers, but our model showed a counterintuitive effect: Because of the masking effect, higher flower density may result in fewer discovered flowers.

To study this effect, we computed the total number of flowers discovered by a bee colony as a function of density and flower size. We considered a field with flowers of a given size uniformly and randomly distributed with a given flower density, simulated 1000 exploration trips, and counted the number of flowers that were discovered at least once. When we performed this simulation neglecting the masking effect (i.e., assuming that a bee discovers all the flowers that intersect with its trajectory, not being affected by such discoveries), we found that the number of flowers discovered increased with flower density and flower size, as these factors make flowers more plentiful and easier to find (Fig. 5, dashed lines). However, the masking effect reverses this trend (Fig. 5, solid lines): For low densities, the masking effect is weak and the number of discovered flowers increases with density, but at high flower densities bees become “trapped” around the nest by the flowers immediately surrounding it, which accumulate most of the visits. Therefore, there is an optimum density that results in the highest number of different flowers discovered. Since the masking effect is stronger for larger flowers, the effect of size also reversed, with the number of discovered flowers decreasing as flower size increases (Fig. 5, solid lines).

**Fig 5.**
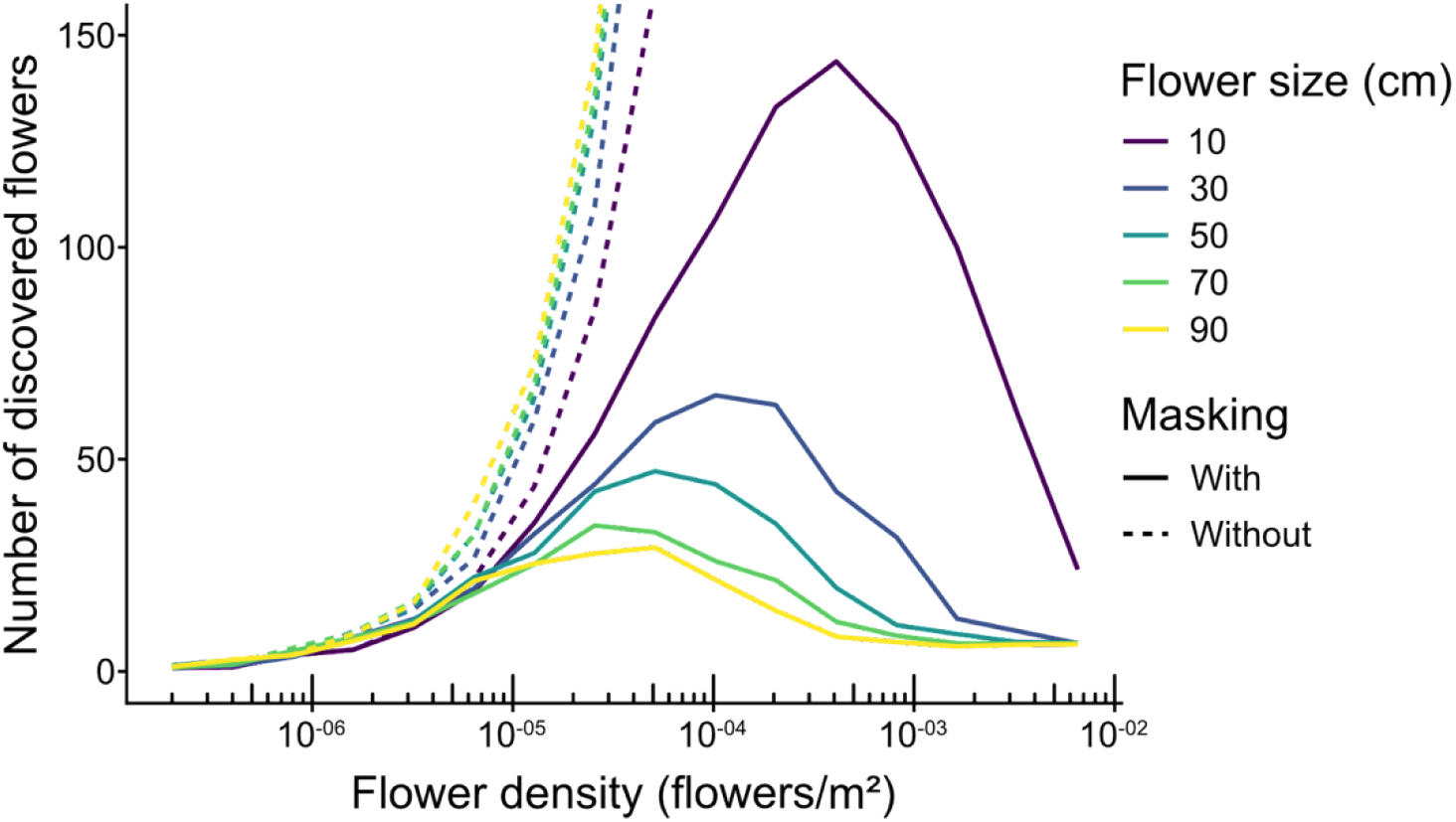
Number of different flowers discovered by a group of bees as a function of flower density. Number of different flowers discovered in 1000 exploration trips of 900 s, in an environment with randomly distributed flowers. Results are averaged over 10 simulations, keeping the environment fixed for every simulation. Solid lines: Probability calculated taking into account the masking effect (i.e., only counting the first flower that was discovered in each trip). Dotted lines: Probability calculated without taking into account the masking effect.

## Discussion

How pollinators search for flowers is of fundamental importance but remains poorly understood. Here we developed a realistic model of bee search movements based on their observed tendency to make exploratory loops that start and end at their nest location. Our model, calibrated with real behavioral data, produces two-dimensional trajectories with progressive changes of direction driven by the continuous evolution of the angular velocity *ω*(*t*). Using this approach, we documented a neglected yet potentially fundamentally important effect for bee foraging success and pollination: a perceptual masking effect that influences the probability of bees to discover flowers depending not only based on their size and spatial location, but also on the presence and characteristics of other flowers around them.

Previous models assume that bees explore the environment randomly using Lévy flights or other diffusive processes (12,21,22). In a diffusive model, individuals are able to wander away from the nest indefinitely if given enough time. In contrast to these models, our model replicates looping trajectories observed in real bees (14,40), which confines the presence of individuals around a nest. As a consequence of the periodic returns of bees to the nest, their distribution becomes independent of the time given to explore. This result has the important consequence that, under the assumptions of our model, longer simulation durations will result in a more thorough exploitation and pollination of the area around the nest, but not in a larger area being exploited and pollinated.

By explicitly simulating individual trajectories in complex environments, our model revealed how the presence of a flower may decrease the probability of discovering another. We named this phenomenon a “perceptual masking effect” and explored its consequences in the probability of bees to discover flowers, ultimately extrapolating on pollination and bee foraging success. At the plant level, flowers distant to the nest were more often visited in low density environments, suggesting lower probabilities of pollination at high densities. Therefore, the area that is pollinated around the nest decreases when the flower density increases. The overall distribution of flower patches directly impacts their pollination and should be taken into account when designing strategies for crop production and assisted pollination. At the bee colony level, insects tended to find more flowers when they were small and at medium densities, suggesting that there is an optimal flower size and density at which collective foraging efficiency is optimized (although the effect of size on foraging efficiency will be compounded with the greater reward provided by bigger flowers on average). The perceptual masking effect may also have an impact on site learning and the formation of traplines: At different scales, flower patches may not be discovered in the same order and the probability of forming an optimal route may depend on the scale at which exploration occurs.

Our search model is a scaffold for future characterization of the movement of bees across time and landscapes. Although we limited our study to flower discovery probability, and therefore only provided predictions for first flower discovery, the model could be used to investigate the full foraging trips of bees, and how they change through time as bees acquire experience with their environment and develop spatial memories (13). It would be particularly interesting to integrate this exploration model into existing learning exploitation models proposed to replicate route formation by bees (20–22). Once a flower is discovered, its location can be learned, and new exploration may start, ultimately allowing for the establishment of traplines. This would be modelled via a modification of the attraction component, which can be modified to point towards previously-discovered flowers instead of the nest. Importantly, model predictions (flower discovery probability, visitation order, flight trajectories) can be experimentally tested and the model calibrated for specific study species. This will facilitate improvement and validation for potential applications. For instance, robust predictive models of bee movements including both exploration and exploitation would be particularly useful for improving precision pollination (to maximize crop pollination), pollinator conservation (to ensure population growth and maintenance), but also in ecotoxicology (to avoid exposure of bees to agrochemicals) and legislation (to avoid unwanted gene flow between plants). Beyond pollinators, our minimal persistent turning walker model could be calibrated to apply to a wide range of species, providing a scaffold for further exploration of the broader interactions between central place foraging animals and their environment.

More broadly speaking, the Persistent Turning Walker model has inspired some developments in other animals, especially fish (41–43) as well as in robotics (44) where is has been proven to display better coverage properties than classical random walks(45). The addition of the attraction component paves the avenue for further developments of this model in these areas as well.

## Methods

The codes used to perform all the simulations, data analyses and figures are available in the Supplementary Information (Zip file S3).

### Modeling nest and flower detection

Bumblebees can detect an object when it forms an angle of 3° on the retina of their compound eyes (46). Therefore, for every model simulation, we set the flowers’ size and calculated the distance at which the bees are able to detect them. We call this the “perception distance”. We considered that a bee visited a flower when it was located at a distance to the bee inferior to the perception distance. We did not take into account the olfactory perception since it could be less reliable because of other factors like wind direction and the flower’s species. However, if taken into account, it would only impact the perception distance of the flowers and the results would not be qualitatively different.

### Analysis of experimental data

We used the dataset of Pasquaretta et al. (39) in which the authors tracked exploratory flight trajectories of bumblebees in the field with a harmonic radar. Bees carrying a transponder were released from a colony nest box located in the middle of a large and flat open field, and performed exploration flights without any spatial limitation. The radar recorded the location of the bees every 3.3s over a distance of ca. 800 m, and with an accuracy of approximately 2 m (14). The bees were tested until they found one of three 20-cm artificial flowers randomly scattered in the field. The position of these flowers was changed whenever one of them was found to prevent the bees from learning their location, but their presence may still affect the bees’ trajectories. We first attempted to control for this factor by removing all trajectories where bees passed near an artificial flower, but this introduced a significant bias towards short trajectories, because bees are less likely to find a flower when they stay near the nest. Therefore, we used the full dataset, and in order to remove the effect of the bees hovering around and exploiting the artificial flowers, we summarized all the points detected in an area within 6 m of an artificial flower as a single point at the location of the flower. This threshold of 6 m was derived from a 4 m perception distance corresponding to 20 cm flowers, plus 2 m to account for the experimental noise. All trajectories are given in S2-section V.

### Dividing trajectories into loops

In order to quantify the trajectories, we divided the tracks into “loops”. We defined a loop as a fragment of a trajectory that starts when the bee leaves that nest and finishes when it enters back. The colony nest box used in the experiments was rectangular, with a diagonal of 37 cm, meaning that the bees were able to see it at approximately 7m. However, in this case we set a higher threshold of 13 m to avoid including learning flights (i.e., flights during which the bee makes characteristic loops to acquire visual memories of target locations such as the nest for navigation (38) into the set of exploratory data. While our model does not produce learning flights, for consistency we also used the 13-m radius around the nest in our simulations.

### Model simulations

All simulations start at the nest (which is located at position 0,0), with a random initial direction, and with zero angular velocity. To simulate the trajectories, we discretized the model using the on a time step *△t* = 0.01. Therefore, at every step we calculated the direction θ(*t* + Δ*t*) as

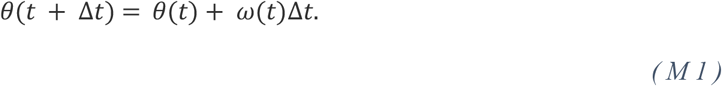

We then calculated the velocity *v*(*t* + Δ*t*) with

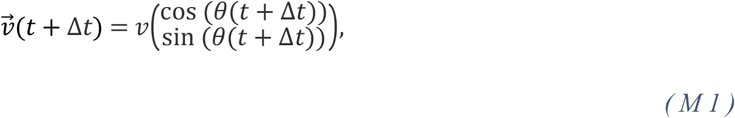

where *v* is the speed, which is a constant in our model. Then, the new position is

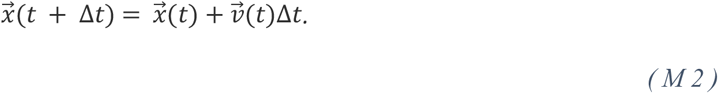

Lastly, we calculated the angular speed *ω*(*t* + Δ*t*). For this, we used the Green functions for Ornstein-Uhlenbeck processes over Δ*t* (see (30) for details), obtaining

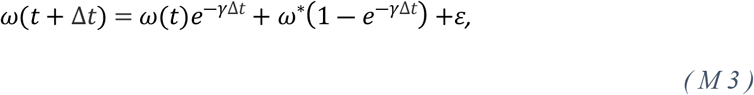

where *ω*\* is the target angular speed (governed by Equations 2 and 3), and *ε* is a random number governed by a Gaussian distribution with mean 0 and variance

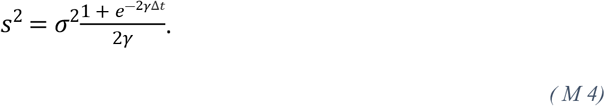

### Parameter fitting

In order to fit the parameters of the model, we explored systematically all relevant combinations within the relevant range for each parameter. To do this more efficiently, we substituted the variance of the noise introduced by the Wiener’s process (σ) for the variance of ω, which has a more direct impact on the experimental data. These two variables are related by (30):

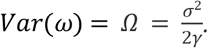

We also defined

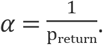

We simulated 10^3^ loops using 6160 different combinations of the parameters of the model (*γ,Ω,α,η*\*)

- *γ* ∈ (0.5,0.6,0.7,0.8,0.9,1.0,1.1,1.2,1.3,1.4,1.5)
- *Ω*∈ (0.01,0.03,0.05,0.06,0.07,0.08,0.09,0.1,0.125,0.15)
- *α* ∈ (10,20,25,30,35,40,50)
- *η*\* ∈ (0.05,0.1,0.15,0.2,0.25,0.3,0.35,0.4)

Since the four observables are heterogeneous (two are continuous measures, two are discrete), a cost function averaging over the four had to be designed to ensure that each observable is given the same weight. For each observable, we computed one score as a distance from the simulated set to the observed set. For the two continuous observables (loop length and extension), the score was computed as the area between the observed cumulative distribution function, and the simulated one. For the two discrete observables (numbers of self-intersection and re-departures), the score was computed, as the absolute difference between the two probability distributions. This yielded four distributions of scores over the 6160 combinations. Scores were then translated into their quantile their corresponding cumulative distribution (e.g., a score translated into 0.12 means that it is within the lowest 12%). Finally, we retained the combination that yielded the lower quantile averaged over the four observables (see S2, section 2).

The best parameters combination was found to be: *γ* = 1.0 *s*^-1^, *Ω* = 0.07 rad^2^/s^-2^, *α* = 30 *s* and *η\** = 0.2 *s*^-1^. It corresponds to the marginal local minima for the four observables (see S2, section 2). Simulated trajectories closely resemble data trajectories (Fig 1B), and the model is able to produce loops with an elongated shape, as well as a diversity of loop lengths.

This unique set of parameters assumes that all bees are identical, while in reality inter-individual differences exist (Fig. S1), for example due to differences in age, experience, learning or size (47,48). However, each bee can display a large diversity of loop parameters, covering a similar range as the overall population (Fig SI-1). We therefore considered that separate fits for each individual were not justified. The fact that our model reproduces not only the mean but also the variability of the four observables we defined (Fig. 3) supports this choice.

## Acknowledgments

We thank Thibault Dubois, Tamara Gómez Moracho, Cristian Pasquaretta, Joe Woodgate, James Makinson, Joanna Brebner and Lars Chittka for sharing their data of bumblebee flight tracks using radar.

## Funding

AM was supported by a PhD Fellowship from the French Government. ML was supported by grants of the Agence Nationale de la Recherche (3DNaviBee ANR-19-CE37-0024), and the European Commission (FEDER ECONECT MP0021763, ERC Cog BEE-MOVE GA101002644). APE acknowledges funding from a CNRS Momentum grant (https://www.cnrs.fr/) and a Fyssen Foundation Research grant (https://www.fondationfyssen.fr/en/).

The funders had no role in study design, data collection and analysis, decision to publish, or preparation of the manuscript.

## Supporting information

**S1 Fig. Variability of each observable across individuals in the experimental dataset. (A)** Loop lengths (m) for each bee, as defined in Fig. 3 in the main text. Boxplots, show the median (middle line), 25 and 75% quantiles (box), range of data within 1.5 interquartile deviations (whiskers), and outliers (dots). **(B)** Same as A but for the loop extension (maximum distance between the nest and the individual). **(C)** Same as A, but for the number of re-departures per 100m traveled. A re-departure is defined as three consecutive positions such that the second position is closer to the nest than the first one, but the third is again further away than the second. **(D)** Same as A but for the intersections (number of times the loop intersects with itself)

**S2 Text. Raw results and figures**

**S3 Data and code sources for analysis and simulations.**

## Notes

### Competing Interest Statement

The authors have declared no competing interest.

